# Effects of regulatory network organization and environment on PmrD connector activity and polymyxin resistance in *Klebsiella pneumoniae* and *Escherichia coli*

**DOI:** 10.1101/2020.05.05.080051

**Authors:** Annie I. Chen, Jun Zhu, Mark Goulian

## Abstract

Polymyxins are a class of cyclic peptides with antimicrobial activity against Gram-negative bacteria. Resistance to these compounds is often mediated by pathways that also confer resistance to host antimicrobial peptides. In *Enterobacteriaceae*, the PhoQ/PhoP and PmrB/PmrA two-component systems regulate many of the genes associated with these resistance mechanisms. In *K. pneumoniae*, spontaneous polymyxin resistance is frequently acquired through inactivation of the gene *mgrB*, which encodes a negative regulator of PhoQ. However, this resistance mechanism has not been reported in other genera of *Enterobacteriaceae*, despite the presence of *mgrB* among many members of this family. In addition, the frequency of developing spontaneous resistance to the antimicrobial peptide polymyxin through chromosomal mutations is much higher in *Klebsiella* compared to *Salmonella* or *E. coli*. Here we show that in *K. pneumoniae*, PmrD is not required for polymyxin resistance arising from inactivation of *mgrB*. In addition, we show that in *E. coli*, the protein PmrD can activate PmrA under certain conditions. Our results suggest that the importance of PmrD connector activity in polymyxin resistance depends on both the network organization and on the environmental conditions associated with PmrB stimulation.

## Introduction

Polymyxins, such as colistin and polymyxin B, have emerged as last-resort antibiotics for many multidrug-resistant Gram-negative bacteria (1). These compounds are cationic antimicrobial peptides that are thought to disrupt the bacterial outer membrane through interactions with negatively charged lipopolysaccharides (LPS) (2–6). In addition to their role in treating infections, polymyxins have also been used in the laboratory as a tool to study pathways in bacteria that mediate resistance to host antimicrobial peptides. Several mechanisms of polymyxin resistance have been identified among Gram-negative bacteria, with the most common mechanism involving modification of the LPS (7, 8).

In many *Enterobacteriaceae*, polymyxin resistance is controlled primarily by the PmrB/PmrA and PhoQ/PhoP two-component systems (9–11). The PmrB/PmrA system regulates the expression of enzymes that covalently modify the lipid A to decrease the negative charge of the outer membrane (9, 12, 13). The sensor kinase PmrB has both kinase and phosphatase activities; activation of PmrB through ferric iron or mild acidity leads to increased levels of phosphorylated PmrA (14, 15). Phosphorylated PmrA activates expression of several genes that encode LPS modification enzymes, including *ugd* and the *arnBCADTEF* operon, whose protein products mediate addition of 4-aminoarabinose to lipid A, and *pmrC*, which encodes an enzyme that modifies the lipid A with phosphoethanolamine (16, 17). PmrA phosphorylation can also be regulated indirectly by the PhoQ/PhoP two-component system (18), which is activated by low magnesium and cationic antimicrobial peptides (19–21). PhoP regulates expression of the gene *pmrD*, which encodes a small protein that inhibits dephosphorylation of PmrA by PmrB (22–25). PmrD is an example of a “connector” protein (because it connects the PhoQ/PhoP and PmrB/PmrA systems) (26), and the indirect regulation of PmrA by PhoP via PmrD is referred to as the connector-mediated pathway. In *Klebsiella*, the PhoQ/PhoP system regulates *arnBCADTEF* not only indirectly via PmrD but also through a second mechanism, namely direct activation of the *arnB* promoter by PhoP (27). This direct activation is absent in *Salmonella* and *E. coli* (Fig. 1).

**Figure 1.**
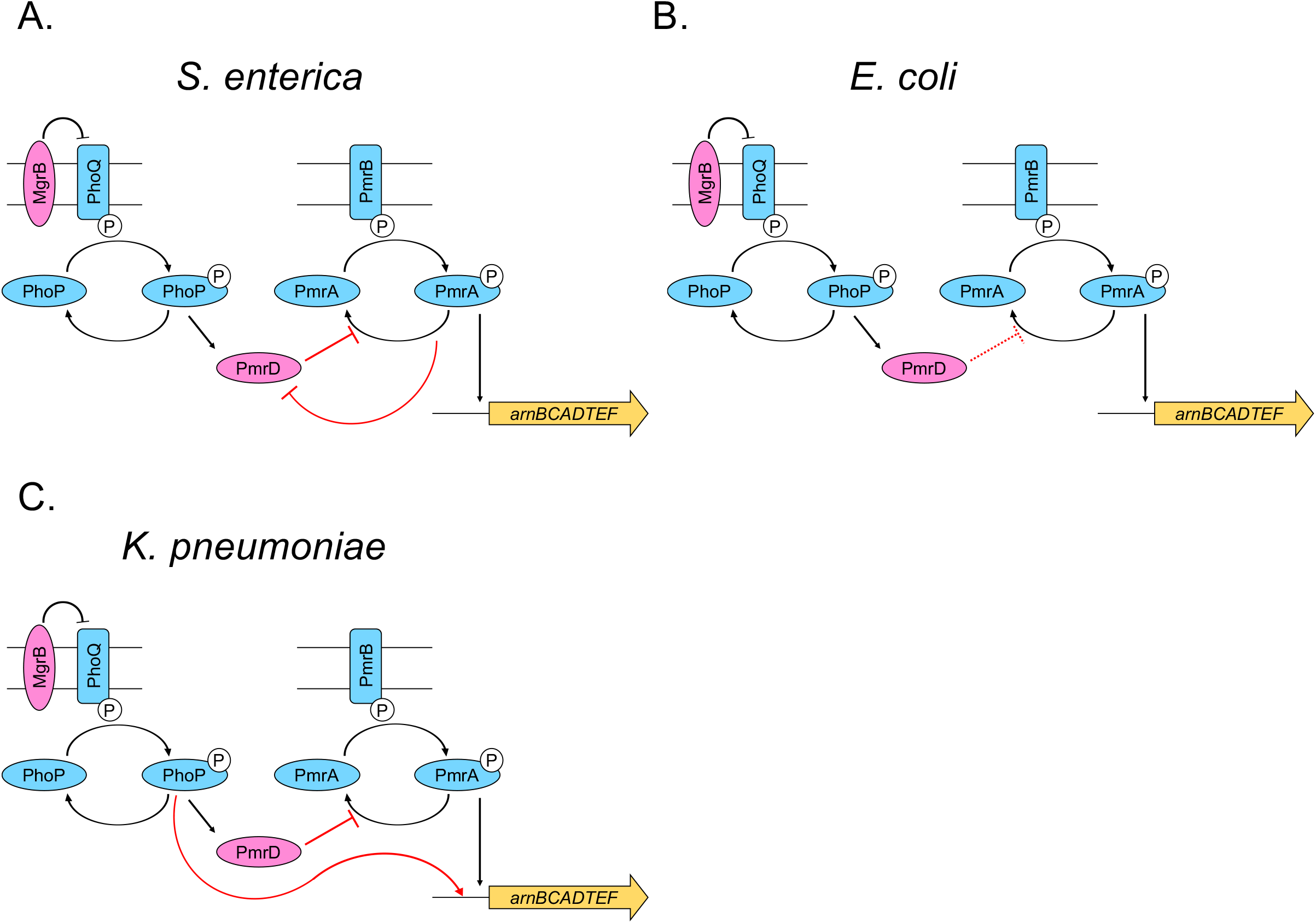
Schematic of polymyxin resistance network in different genera of *Enterobacteriaceae*. **A.** In *S. enterica*, PmrD functions as a connector between PhoQ/PhoP and PmrB/PmrA, and PmrA also negatively regulates transcription of PmrD (24, 59). **B.** Previous studies did not detect PmrD connector activity in *E. coli* (41). Since in this study we report that PmrD can function as a connector in *E. coli* under certain conditions, we indicate the connector activity with a dotted line. **C.** In *K. pneumoniae*, PmrD can function as a connector, and PhoP can also directly regulate transcription of the *arnB* operon (27).

Cross-regulation between other two-component systems has also been shown to mediate polymyxin resistance. In some uropathogenic *E. coli*, for example, PmrB can phosphorylate a noncognate response regulator, QseB, which has been shown to play a role in polymyxin resistance (28). In addition, some *K. pneumoniae* strains encode the CrrB/CrrA two-component system, which can activate PmrB through the CrrA-regulated gene *crrC* (29, 30).

Analysis of polymyxin-resistant isolates of *Enterobacteriaceae* in clinical and laboratory settings have identified several mutations that confer polymyxin resistance (7, 31). Curiously, the chromosomal mutations identified in polymyxin-resistant isolates vary among *Enterobacteriaceae*, such as *Klebsiella pneumoniae*, *Salmonella enterica*, and *Escherichia coli*, even though the proteins involved in regulating LPS modifications are largely conserved. In *K. pneumoniae*, mutations in *phoQ*, *phoP*, *pmrB*, and *pmrA* that result in constitutive expression of LPS modifications have been shown to confer polymyxin resistance. Yet the most common mechanism of polymyxin resistance in *K. pneumoniae* is inactivation of the gene *mgrB*, which encodes a negative regulator of PhoQ (32–34). Numerous mutations in *mgrB* have been identified among polymyxin-resistant isolates, including inactivation through insertion sequences, nonsense mutations, or missense mutations. The high frequency of this mechanism of resistance likely reflects the fact that there are many more ways to inactivate a gene compared to the limited number of mutations in a sensor kinase or response regulator gene that lead to constitutive activation. Since mutations in the polymyxin resistance network, either in *mgrB* or in two-component system proteins, all lead to upregulation of LPS modifications, one might expect to find *mgrB* inactivation as a dominant mechanism of resistance in related bacteria. However, inactivation of *mgrB* has not been observed as a mechanism of resistance among other *Enterobacteriaceae*, even though many other genera have *mgrB* orthologs (34).

In contrast to *K. pneumoniae, Salmonella* primarily acquires polymyxin resistance through mutations in the sensor kinase or response regulator of the PmrB/PmrA or PhoQ/PhoP two-component systems (10, 16, 35–37). In *E. coli*, on the other hand, the PmrB/PmrA system alone is the primary target for mutations conferring polymyxin resistance (38–40). Previous reports have shown that stimulation of PhoQ in *E. coli* does not lead to PmrA activation, leading to the conclusion that *E. coli* PmrD cannot function as a connector protein between PhoQ/PhoP and PmrB/PmrA (41). However, the *E. coli* PmrD protein can function as a connector in *Salmonella*, and the *Salmonella* PmrD protein can similarly function as a connector in *E. coli* (41, 42). This difference between the activities of *E. coli* and *Salmonella* PmrD orthologs is apparently due to differences in their ability to inhibit the phosphatase activity of *E. coli* PmrB, which is higher than that of *Salmonella* PmrB (42). These observations raise the question of why *E. coli* has a PmrD protein. However, one study reported an increase in LPS modifications under PhoQ-stimulating conditions in *E. coli* that was dependent on PmrD (43).

Here we investigate the regulation of the *arnB* operon by the PhoQ/PhoP two-component system in *K. pneumoniae* and in *E. coli*. We find that the direct regulation of *arnB* by PhoP in *K. pneumoniae* is critical for *mgrB* inactivation to produce high-level polymyxin resistance. We also find that PmrD can function as a connector between PhoQ/PhoP and PmrB/PmrA in *E. coli* and further show that the role of PmrD as a connector protein depends on the level of PmrB stimulation both in *K. pneumoniae* and in *E. coli*. These results indicate that environmental conditions affect the level of cross-regulation between PhoQ/PhoP and PmrB/PmrA.

## Results

### Frequency of spontaneous polymyxin resistance is higher in *K. pneumoniae* than in *Salmonella* or *E. coli* and is consistent with differences in the effect of *mgrB* inactivation

To test whether deletion of *mgrB* could improve polymyxin resistance in other genera of *Enterobacteriaceae* in addition to *Klebsiella*, we compared the minimum inhibitory concentration (MIC) of polymyxin B for wild-type and Δ*mgrB* strains of *K. pneumoniae* MGH 78578, *S. enterica* serovar Typhimurium (*S.* Typhimurium) 14028s, and *E. coli* MG1655. As expected, we found that the MIC of polymyxin B was much higher (~16-32 times higher) for a *K. pneumoniae* Δ*mgrB* strain compared to wild type (Table 1). In contrast, deletion of *mgrB* in *Salmonella* only increased the MIC by twofold (Table 1), even though activation of PhoQ through low magnesium increases polymyxin tolerance in *Salmonella* (44). Deletion of *mgrB* did not confer any detectable protection against polymyxin in *E. coli* under the conditions of the assay, which is consistent with previous reports that activation of PhoQ does not lead to activation of PmrA in *E. coli* (41).

**Table 1.**
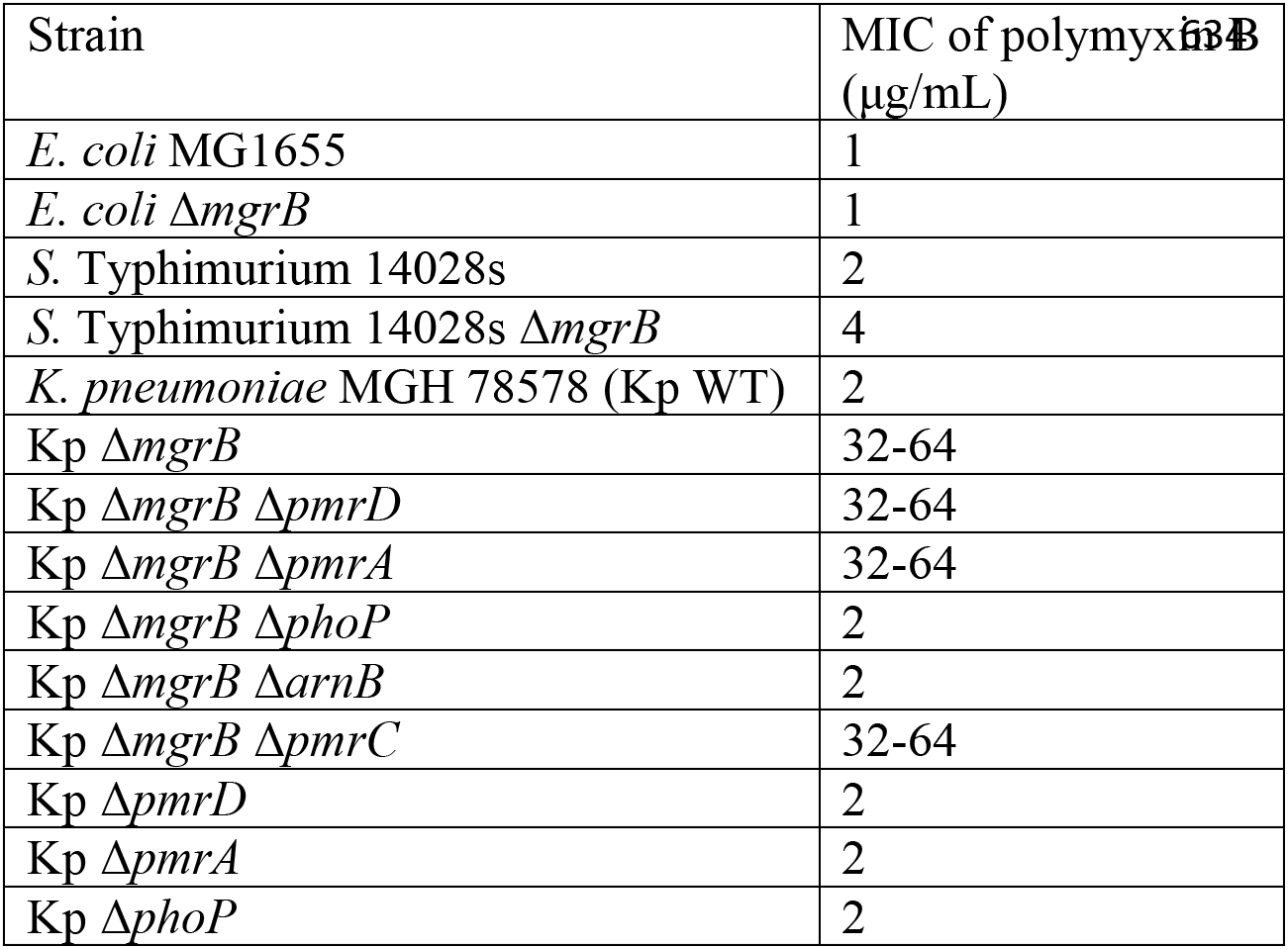
MIC of polymyxin B (μg/mL)

The above observations suggest that the frequency of spontaneous polymyxin resistance would be higher for *K. pneumoniae* than for *Salmonella* and *E. coli*, since inactivating mutations of *mgrB* are likely to arise with higher frequency than the change-of-function mutations associated with other mechanisms of polymyxin B resistance. To test this hypothesis, we plated overnight cultures of wild-type *K. pneumoniae*, *S.* Typhimurium, and *E. coli* on LB agar containing 10 μg/mL polymyxin B to select for spontaneous resistance. We found that indeed the frequency of spontaneous polymyxin resistance was much higher for *K. pneumoniae* (Fig. 2). We also tested eight colonies at random and found that all had inactivating mutations in *mgrB* (7 insertion sequences and one nonsense mutation), indicating that inactivation of *mgrB* accounts for the high frequency of spontaneous polymyxin resistance in *K. pneumoniae*.

**Figure 2.**
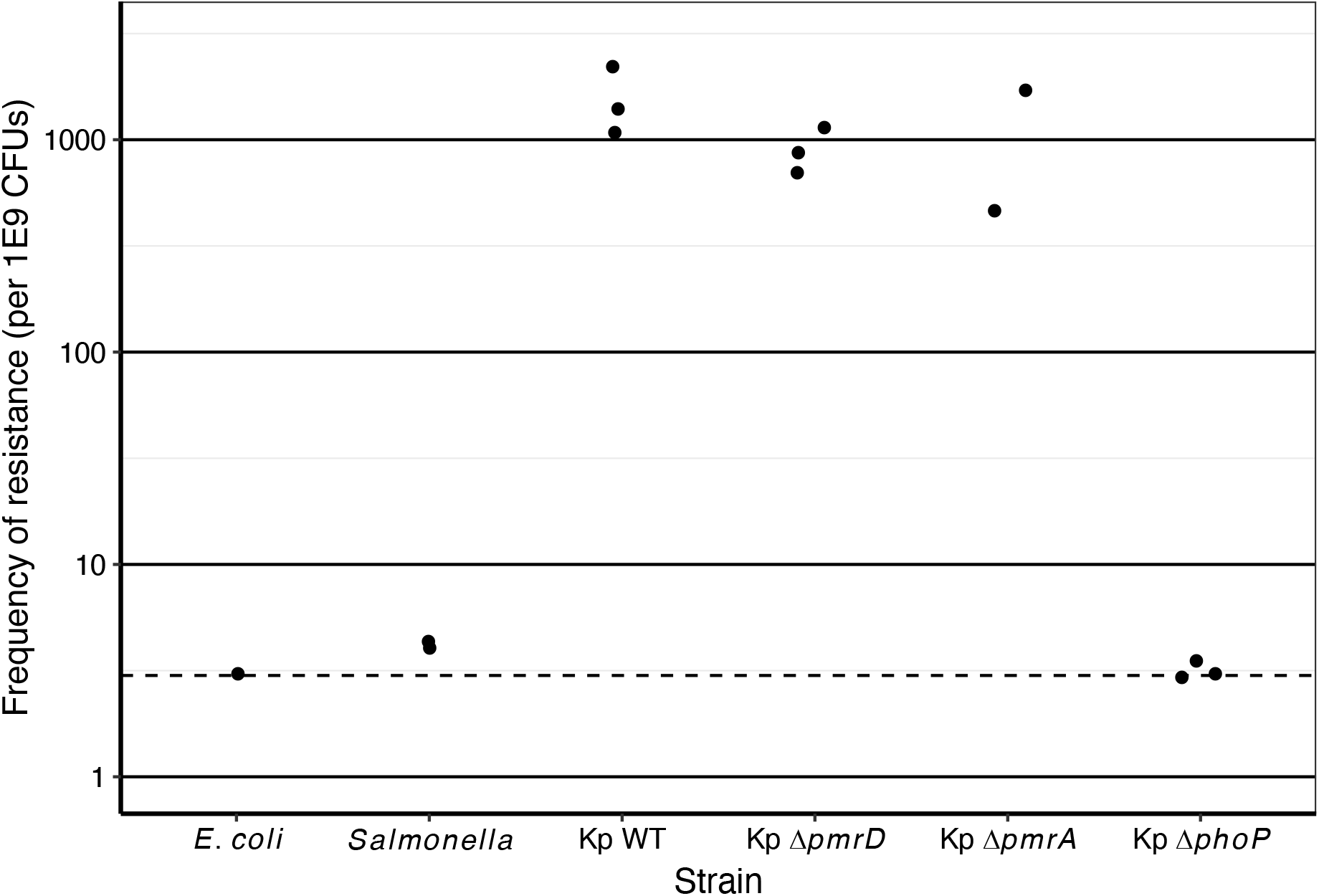
Frequency of spontaneous polymyxin resistance. Cells were grown in LB and plated on LB with 10 μg/mL polymyxin B. Dilutions of the culture were also plated on LB. The frequency of spontaneous polymyxin resistance was determined by dividing the number of resistant CFUs (colony forming units) on the LB + polymyxin plate by the number of CFUs on the LB plate. Each dot represents a biological replicate. The dotted line denotes the limit of detection. Strains used: MGH 78578, ACK25, ACK24, ACK26, 14028s, MG1655.

### PmrD and PmrA play a minor role in polymyxin resistance in a *K. pneumoniae* Δ*mgrB* strain

In *Klebsiella*, PhoP directly activates transcription of the *arnB* operon, a mode of *arnB* regulation that is not found in *Salmonella* or *E. coli* (Fig. 1) (27). We therefore tested the relative importance of direct regulation and of the connector-mediated pathway in polymyxin resistance in *K. pneumoniae*. We found that deletion of *pmrD* in a Δ*mgrB* strain had no effect on the polymyxin B MIC, suggesting that the connector-mediated pathway between PhoQ/PhoP and PmrB/PmrA was not necessary for resistance in this bacterium (Table 1). In addition, deletion of *pmrA* similarly did not affect the MIC of polymyxin B, which is consistent with previous observations (45). On the other hand, resistance was suppressed by deleting *phoP*, indicating that PhoP is necessary for the polymyxin resistance of a Δ*mgrB* strain. To detect potentially smaller effects of polymyxin among strains, we also used a polymyxin B sensitivity assay in which exponential-phase cells were treated with polymyxin for one hour, and then plated to determine the survival relative to untreated controls (see Materials and Methods). We found that deleting either *pmrD* or *pmrA* in a Δ*mgrB* strain had little effect on survival, whereas deleting *phoP* suppressed polymyxin tolerance of a Δ*mgrB* strain (Fig. 3). We also found that the frequency of spontaneous polymyxin resistance was similar for *K. pneumoniae* wild-type, Δ*pmrD*, and Δ*pmrA* strains, and was much higher than that of a Δ*phoP* strain (Fig. 2), consistent with the above observations. Taken together, the above results indicate that polymyxin resistance mediated by inactivation of *mgrB* requires PhoP but does not require PmrD or PmrA.

**Figure 3.**
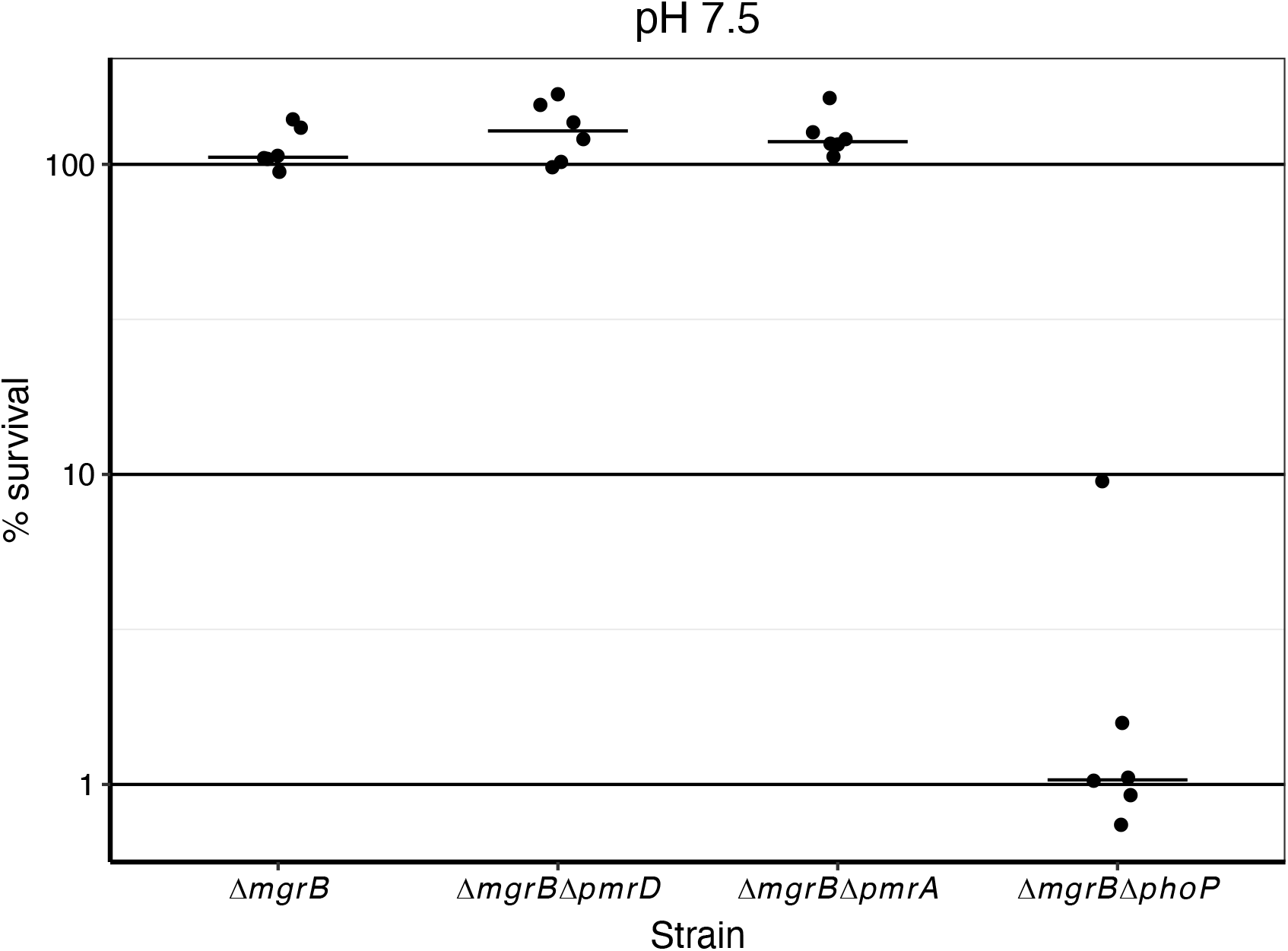
Relative roles of PmrD, PmrA, and PhoP in polymyxin tolerance of a *K. pneumoniae* Δ*mgrB* strain. Strains were grown in N-minimal medium pH 7.5 with 1 mM MgCl_2_ to exponential phase and diluted into PBS + 1 mM MgCl_2_ with or without 1 μg/mL polymyxin B and incubated for 1 hour before plating. Percent survival was calculated based on CFUs after treatment with polymyxin relative to those of untreated samples. In some cases, the calculated survival may have been over 100% due to pipetting error or the completion of a round of replication during the killing assay. Horizontal lines represent the medians of the pooled data from two independent experiments, and each dot represents a biological replicate. Strains used: ACK9, ACK21, ACK20, and ACK28.

To assess the importance of aminoarabinose and phosphoethanolamine lipid A modification pathways to polymyxin resistance in a *K. pneumoniae* Δ*mgrB* strain, we tested the effect of deleting either *arnB* or *pmrC*. Deletion of *pmrC* did not affect polymyxin resistance. On the other hand, deletion of *arnB* in a Δ*mgrB* strain lowered the MIC to that of wild type, suggesting that the *arnB* gene is critical for polymyxin resistance mediated by *mgrB* inactivation (Table 1), which is consistent with the results of a previous study (45).

### PmrD connector activity in *K. pneumoniae* depends on environmental stimuli

In the assays described above, PmrD did not affect polymyxin sensitivity of a *K. pneumoniae* Δ*mgrB* strain. However, a previous study found that deletion of *pmrD* in wild-type *K. pneumoniae* affects polymyxin tolerance (25). We therefore considered the possibility that these different observations reflect differences in growth conditions. We reasoned that if either PhoQ or PmrB is sufficiently activated, then PmrD may not be important for polymyxin resistance, since both PhoP and PmrA can directly activate the *arnB* operon. However, for low or intermediate PmrB and PhoQ stimulation, PmrD may play an important role in limiting PmrA dephosphorylation and promoting polymyxin resistance.

To test this hypothesis, we grew cells in different concentrations of FeSO_4_, which stimulates PmrB (14), and measured polymyxin B susceptibility. In the absence of iron addition, wild-type *K. pneumoniae* cells showed ~1% survival from treatment with 1 μg/mL polymyxin B. When cells were pre-treated with 15 μM FeSO_4_, the survival increased to ~100%, and this increase in survival depended on *pmrD*, *pmrA*, and *phoP* (Fig. 4). When cells were pre-induced with 100 μM FeSO_4_, the survival of Δ*pmrD* cells increased to ~70%, suggesting that high PmrB stimulation decreases the importance of PmrD in promoting polymyxin tolerance. We also found that the survival of a Δ*phoP* strain pre-induced with 100 μM FeSO_4_ was much lower than that of wild type. This observation suggests that, even under the strong PmrB-stimulating conditions of high iron, the direct regulation of the *arnB* operon by PhoP, or regulation of other LPS-modifying genes by PhoP, is critical for survival.

**Figure 4.**
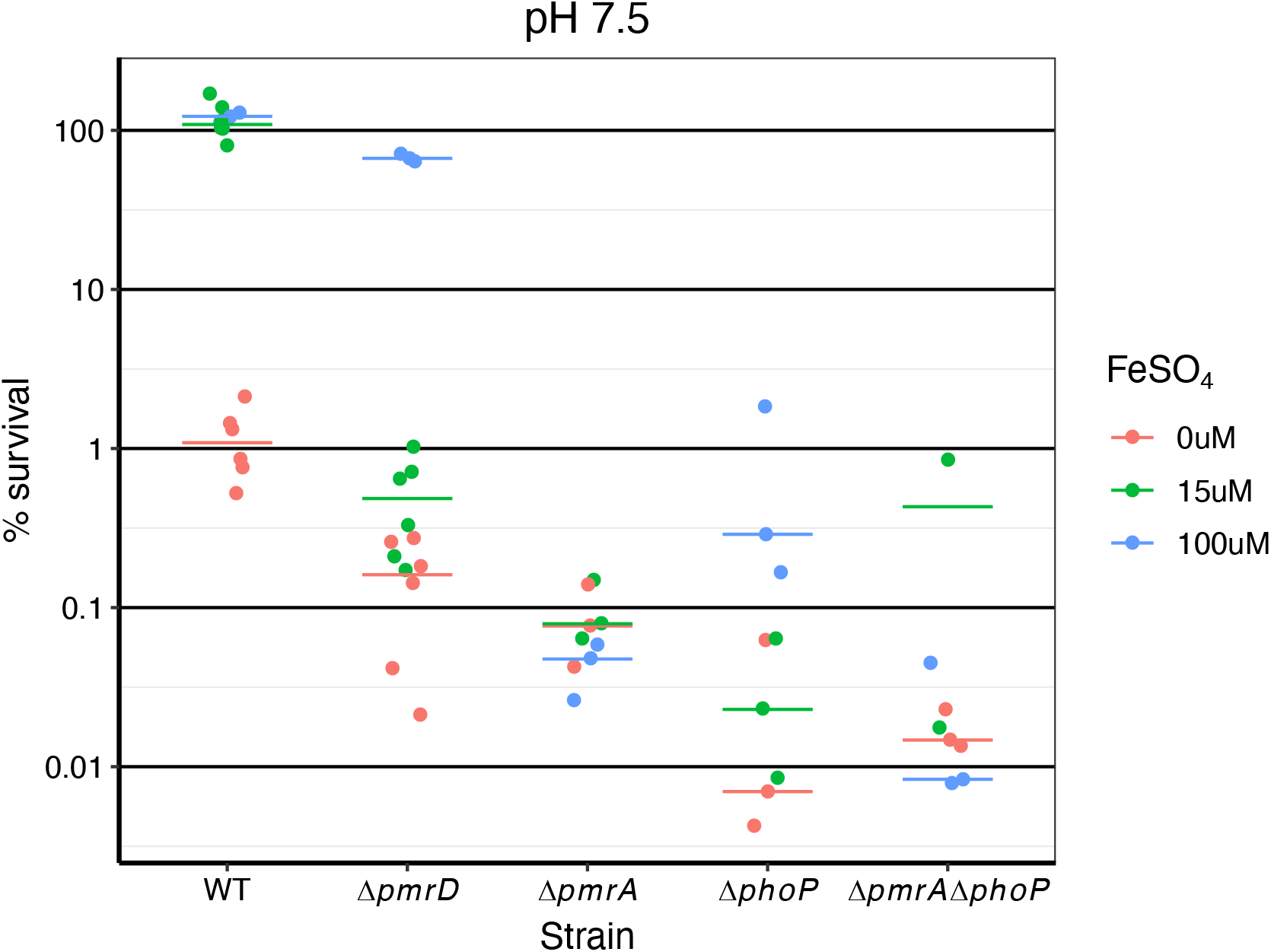
The effect of iron on *K. pneumoniae* polymyxin B tolerance. Polymyxin susceptibility assays were performed as in Figure 3 except that the indicated concentrations of FeSO_4_ were added to the growth medium. Horizontal lines represent the medians of the pooled data from two independent experiments, and each dot represents a biological replicate. Strains used: MGH 78578, ACK25, ACK24, ACK26, and ACK51.

Since mild acidity has been shown to stimulate PhoQ and PmrB in *Salmonella* (15, 46–48), we also tested whether pre-induction with mild acidity could increase polymyxin tolerance in *K. pneumoniae*. We found that wild-type *K. pneumoniae* cells grown in pH 5.8 N-minimal medium showed increased polymyxin tolerance compared to those grown at pH 7.5, and deletion of *pmrD* did not affect survival (Fig. 5). Deletion of *pmrA* showed only a slight decrease in survival compared to that of wild type, suggesting that PhoP may play a more dominant role than PmrA under these conditions.

**Figure 5.**
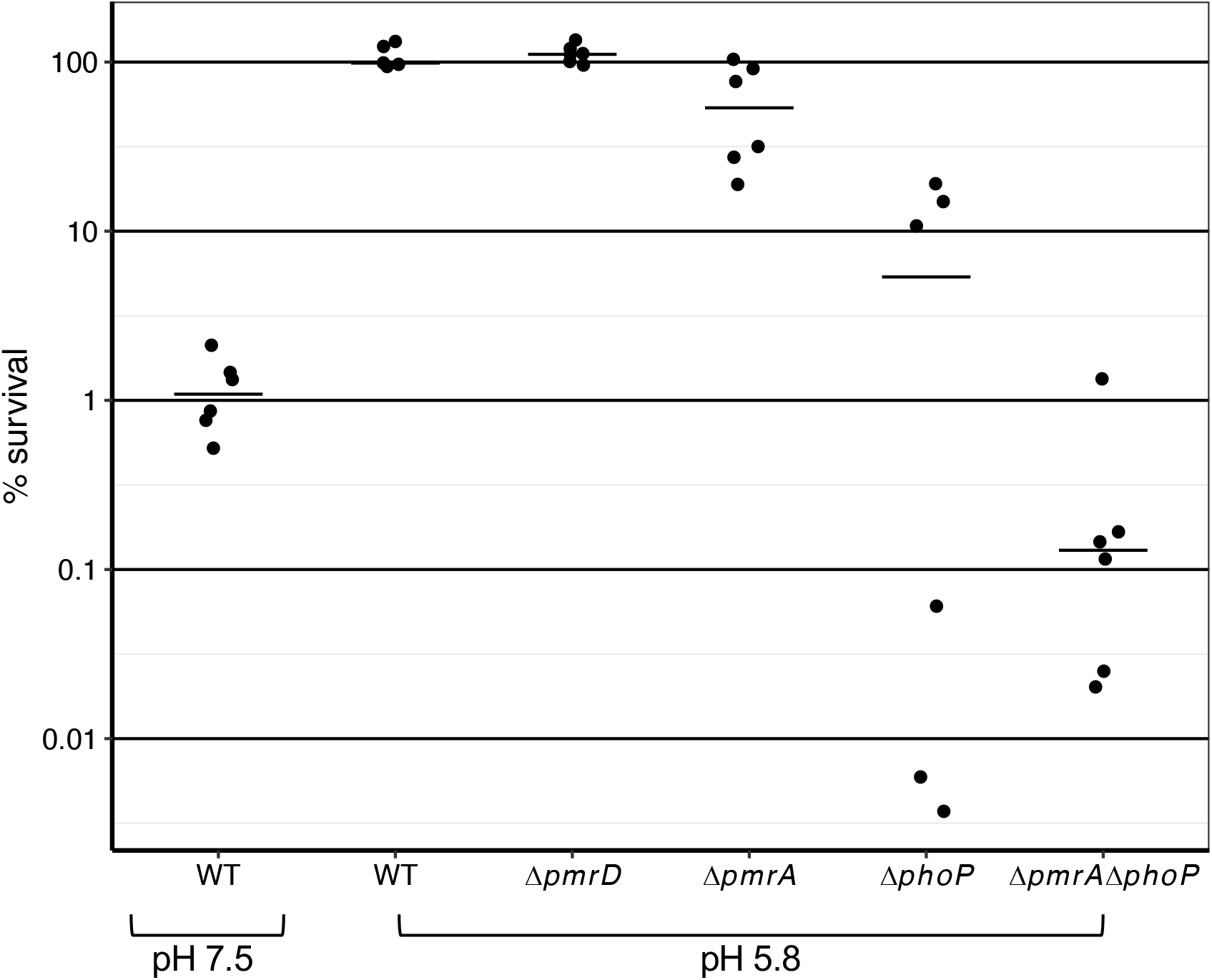
The effect of mild acidity on polymyxin tolerance in *K. pneumoniae*. Cells were grown in N-minimal medium pH 5.8 with 1 mM MgCl_2_ to exponential phase and then polymyxin B assays were performed as described in Figure 3. Horizontal lines represent the medians of the pooled data from two independent experiments, and each dot represents a biological replicate. Strains used: MGH 78578, ACK25, ACK24, ACK26, and ACK51.

Taken together, the above results suggest that PmrD connector activity in *K. pneumoniae* is critical for polymyxin resistance under conditions with relatively low PhoQ stimulation and intermediate PmrB stimulation, such as growth in 1 mM Mg^++^ and 15 μM Fe^++^.

### Iron enables *E. coli* PmrD to function as a connector between PhoQ/PhoP and PmrB/PmrA

In light of our observations that the role of *K. pneumoniae* PmrD in polymyxin resistance depends on environmental conditions, we revisited the role of PmrD in *E. coli.* As discussed above, previous work indicated that *E. coli* PmrD does not function as a connector between PhoQ/PhoP and PmrB/PmrA in *E. coli*, and it was suggested that this lack of connector activity is due to the strong phosphatase activity of *E. coli* PmrB compared, for example, to that of *Salmonella* PmrB (41, 42). However, a recent study reported that *E. coli* PmrD plays a role in polymyxin tolerance when PhoQ is stimulated by low magnesium (43), although PmrD connector activity was not addressed directly. The minimal medium used in the latter study contained 15 μM FeSO_4_. We therefore considered the possibility that the apparent discrepancy between these different studies reflects differences in the level of PmrB stimulation. We reasoned that iron might lower PmrB phosphatase activity and thereby enable PmrD to function as a connector in *E. coli*.

To test this hypothesis, we assayed polymyxin sensitivity of wild-type *E. coli* cells growing in N-minimal medium pH 7.5 with high or low magnesium and with varying concentrations of iron. We first tested the effect of iron supplementation for cells growing in 1 mM Mg^++^, a condition for which *E. coli* PhoQ is only moderately stimulated (49). Addition of FeSO_4_ to either 15 μM or 100 μM did not increase tolerance polymyxin B (Fig. 6A). This result contrasts with the behavior of *K. pneumoniae*, for which 15 μM FeSO_4_ was sufficient to increase survival to ~100% in these growth conditions.

**Figure 6.**
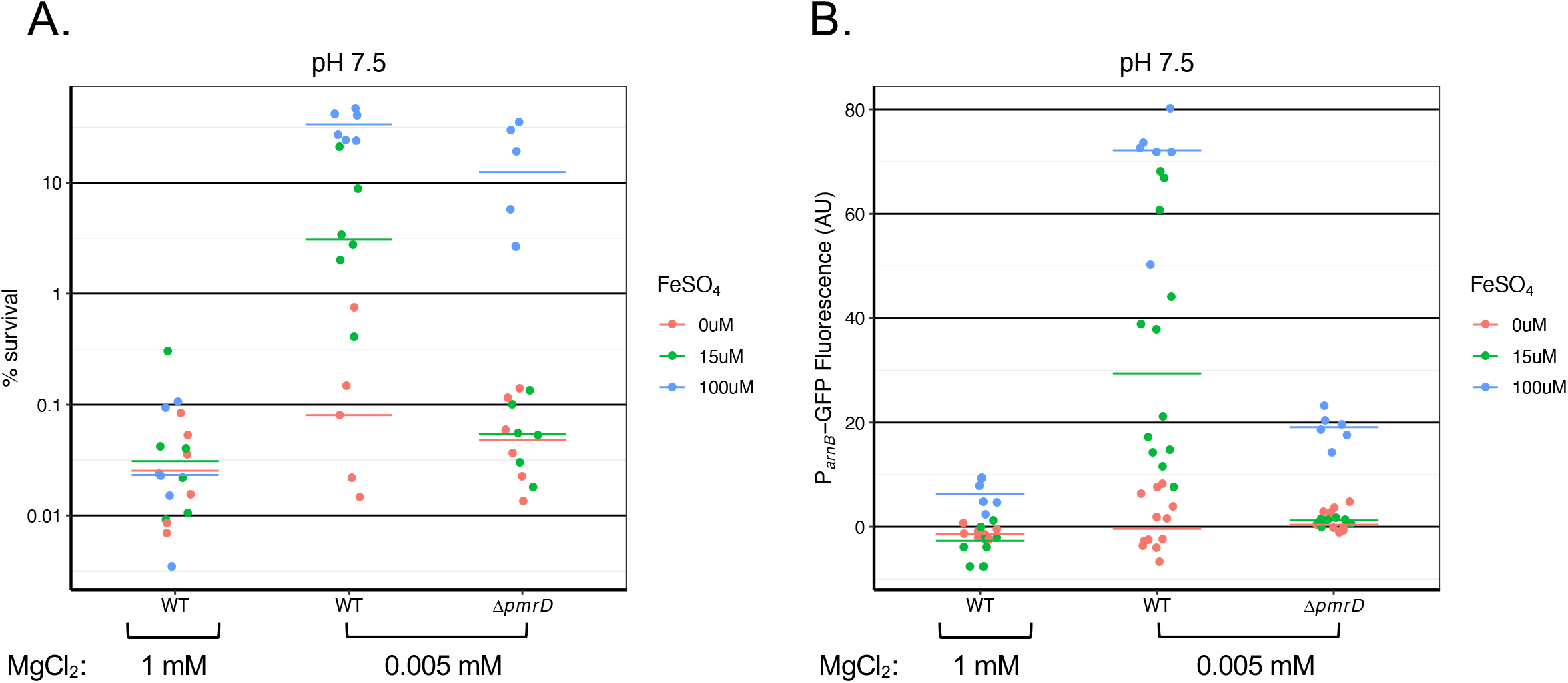
Effect of iron on polymyxin resistance and *arnB* transcription in *E. coli*. **A.** *E. coli* wild-type and Δ*pmrD* strains were grown in N-minimal medium pH 7.5 with the indicated concentrations of MgCl_2_ and FeSO_4_, and polymyxin B assays were then performed as in Figure 3. **B.** Fluorescence of a P_*arnB*_-gfp reporter for cells grown as described in Fig. 6A. Horizontal lines represent the medians of the pooled data from two independent experiments, and each dot represents a biological replicate. Strains used: A) MG1655 and AIC241; B) MG1655/pWKS130, MG1655/pP_*arnB*_-gfp, and AIC241/pP_*arnB*_-gfp.

We next tested cells growing in low magnesium (5 μM Mg^++^), a condition that strongly stimulates PhoQ. Without added iron, low magnesium did not increase polymyxin tolerance relative to 1 mM Mg^++^, consistent with previous reports (41, 42) (Fig. 6A). However, addition of 15 μM FeSO_4_ to the low magnesium medium increased survival more than 10-fold, and addition of 100 μM FeSO_4_ further increased survival. Deletion of *pmrD* abrogated this effect for 15 μM FeSO_4_ but decreased survival only about ~2-fold for 100 μM FeSO_4._ This result suggests that the role of *E. coli* PmrD in polymyxin tolerance is most important when there is strong stimulation of PhoQ and intermediate stimulation of PmrB. In addition, the observation that *pmrD* is less important for polymyxin survival under conditions of low magnesium and 100 μM FeSO_4_ suggests that this high concentration of iron stimulates PmrB sufficiently strongly to increase polymyxin tolerance on its own.

To determine the effect of the above growth conditions on PmrA activity, we used a fluorescent reporter for the PmrA-dependent *arnB* promoter, P_*arnB*_-*gfp*. For wild-type cells growing in high Mg^++^, 100 μM FeSO_4_ produced a relatively small increase in *arnB* transcription (Fig. 6B). Under low magnesium conditions, on the other hand, both 15 μM and 100 μM FeSO_4_ greatly increased *arnB* transcription, consistent with the results of the polymyxin B sensitivity assays. Furthermore, deletion of *pmrD* decreased the effects of iron on *arnB* transcription at low magnesium (Fig. 6B), although the level of GFP fluorescence from Δ*pmrD* cells with 100 μM FeSO_4_ was still higher than that of wild-type cells grown in high magnesium with the same concentration of iron. This latter result is consistent with the observation above that 100 μM iron increases polymyxin tolerance independently of *pmrD* at low magnesium but not at high magnesium and suggests that magnesium exerts both PmrD-dependent and PmrD-independent effects on *E. coli* PmrB/PmrA activity.

### *E. coli* PmrD also functions as a connector at low pH or when PhoQ is strongly stimulated

Since low pH is another PmrB stimulus (in addition to stimulating PhoQ), we tested whether this condition also enables PmrD to function as a connector. We found that growth at pH 5.8 greatly increased GFP fluorescence from P_*arnB*_-*gfp* relative to cells grown at pH 7.5, and this increase depended on *phoP*, *pmrD*, and *pmrA* (Fig. 7). Thus, *E. coli* PmrD connects PhoQ/PhoP with PmrB/PmrA under two conditions that are known to stimulate PmrB.

**Figure 7.**
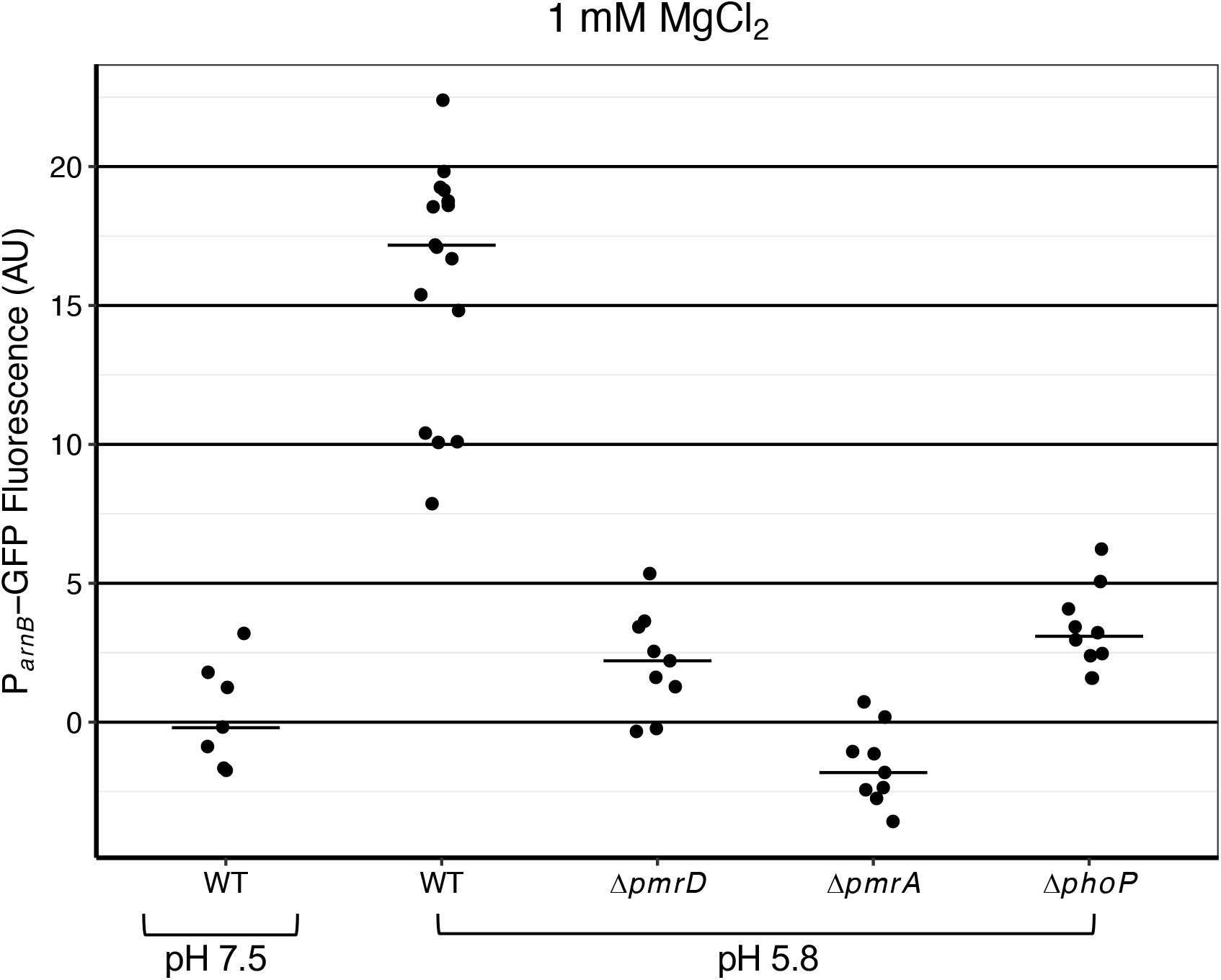
Effect of mild acidity on *arnB* transcription in *E. coli*. *E. coli* were grown in N-minimal medium pH 7.5 or pH 5.8 with 1 mM MgCl_2_ to exponential phase. PmrA activity was measured using a plasmid fluorescent reporter for the *arnB* promoter. Horizontal lines represent the medians of the pooled data from three independent experiments, and each dot represents a biological replicate. Strains used: MG1655/pWKS130, MG1655/pP_*arnB*_-gfp, AIC241/pP_*arnB*_-gfp, AIC204/pP_*arnB*_-gfp, and TIM136/pP_*arnB*_-gfp.

We also wondered whether PmrD could function as a connector even in the absence of iron or low pH if PhoQ were stimulated to a sufficiently high level. We reasoned that strongly stimulated PhoQ/PhoP might produce enough PmrD to effectively overpower the strong phosphatase activity of *E. coli* PmrB. To achieve high levels of PhoQ stimulation, we grew a Δ*mgrB* strain in low magnesium (34, 50). We found that P_*arnB*_-*gfp* transcription increased with decreasing concentrations of magnesium, and this increase in transcription required *pmrD* (Fig. 8). These results suggest that very high stimulation of PhoQ achieved through the combination of low magnesium and deletion of *mgrB* enables PmrD to function as a connector between PhoQ/PhoP and PmrB/PmrA in *E. coli*.

**Figure 8.**
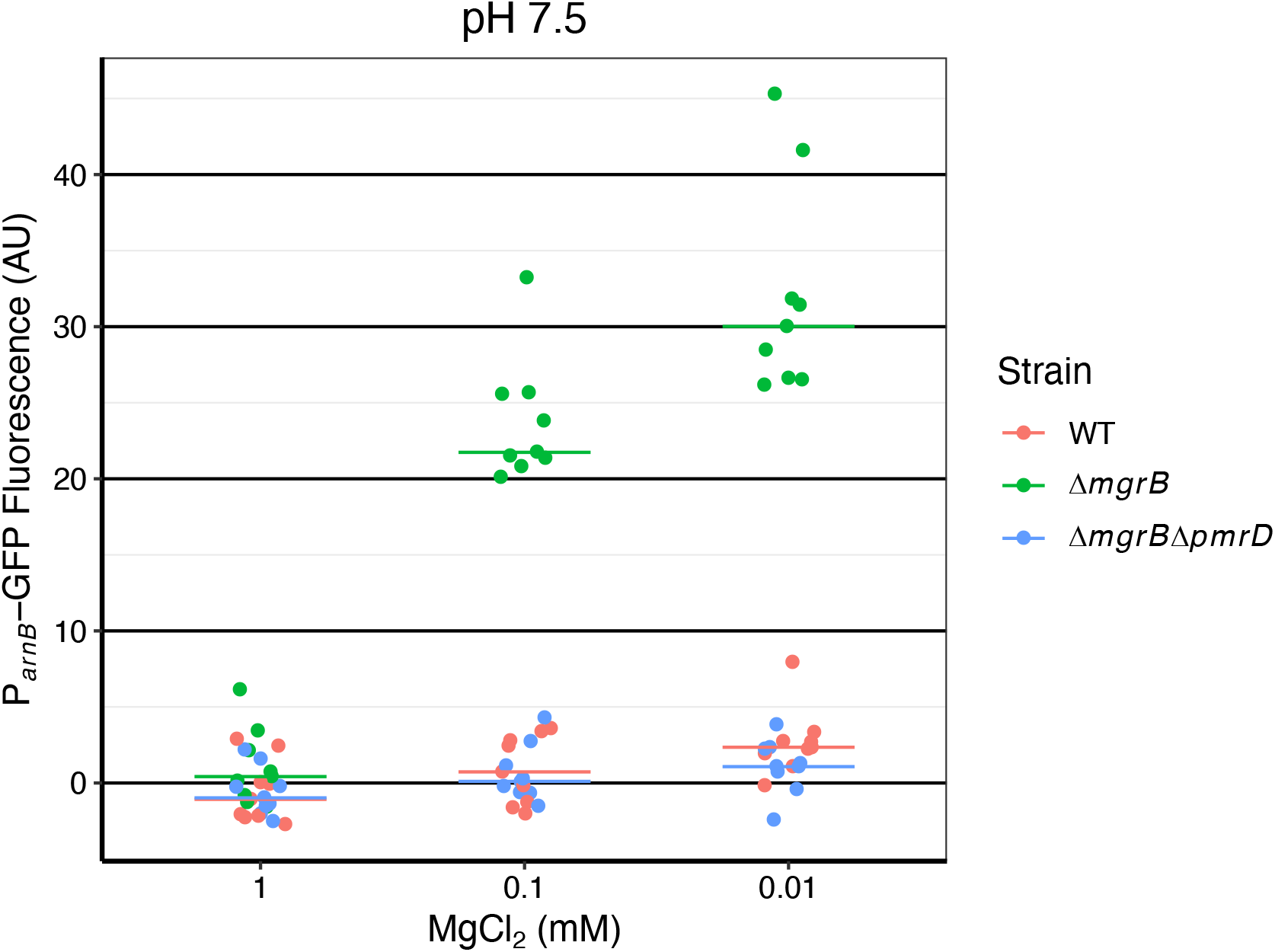
Effect of high PhoQ stimulation on *arnB* transcripton in *E. coli*. Cells were grown in N-minimal medium pH 7.5 with the indicated concentrations of MgCl_2_. PmrA activity was measured using a plasmid fluorescent reporter for the *arnB* promoter. Horizontal lines represent the medians of the pooled data from three independent experiments, and each dot represents a biological replicate. Strains used: MG1655/pWKS130, MG1655/pP_*arnB*_-gfp, MMR155/pP_*arnB*_-gfp, and AIC242/pP_*arnB*_-gfp.

## Discussion

This work extends previous observations concerning natural variation in the organization of the polymyxin resistance network mediated by the PhoQ/PhoP and PmrB/PmrA two-component systems. Our results indicate that differences in the organization of this network account for the fact that *mgrB* inactivation leads to high levels of polymyxin resistance in *Klebsiella* but not in the closely related *Enterobacteriaceae E. coli* and *Salmonella*. Specifically, the direct regulation of the *arnB* operon by PhoP, which occurs in *Klebsiella* but not in *E. coli* and *Salmonella*, is required for this resistance mechanism. This direct regulation is not unique to *Klebsiella* and also appears, for example, in *Yersinia pestis*. It would therefore be interesting to test whether *mgrB* inactivation similarly confers polymyxin resistance in other bacteria with this type of regulation of *arnB*.

The above results also revise our understanding of the polymyxin resistance network in *E. coli*. We find that PmrD functions as a connector between PhoQ/PhoP and PmrB/PmrA in *E. coli*, but this activity depends on appropriate levels of PhoQ and PmrB stimulation. This latter observation explains why *E. coli* connector activity was not observed in previous studies and reconciles results that appeared to be in conflict, but instead likely reflect different levels of PmrB stimulation due to the presence or absence of 15 μM iron in the growth medium (41, 43). As discussed above, we hypothesized that PmrB stimulation lowers the otherwise strong phosphatase activity of *E. coli* PmrB, enabling PmrD to function as a connector. In this sense, our results do not contradict and in fact provide further support for previous observations regarding the differences in the activity of PmrD and PmrB in *E. coli* and *Salmonella* (41, 42). In *Klebsiella*, we also find that PmrD activity depends on the levels of PhoQ and PmrB stimulation, and is most important for polymyxin resistance when PhoQ stimulation is low and PmrB stimulation is intermediate.

For growth in mild acidity, we found that a *K. pneumoniae* Δ*phoP* strain was more sensitive to polymyxin than a Δ*pmrA* strain. We also found that under high iron conditions, deletion of *phoP* showed a significant decrease in survival compared to wild type, even though survival was higher than that of a Δ*phoP* strain grown without added iron. This latter result may indicate that the direct regulation of *arnB* by PhoP contributes to polymyxin resistance even under the strong PmrB-stimulating conditions of high iron. It may also indicate that, despite high PmrB stimulation, additional LPS modification genes regulated by PhoP are needed for full protection against polymyxin (45). Overall, at least under the conditions tested here, PhoP appears to play a dominant role in polymyxin resistance in *Klebsiella*.

The regulatory architecture in *K. pneumoniae*, which consists of direct activation and indirect activation of *arnB* through a connector protein, is an example of a feedforward connector loop. This network topology is characterized by fast activation and slow deactivation of transcription, properties that may be important for bacterial fitness (27, 51, 52). Our results highlight another feature of this architecture: the ability to tune the output in response to multiple environmental signals that act at different points in the network—in this case, signals that stimulate PhoQ (low magnesium), PmrB (iron), or both (pH). Thus, this architecture enables complex environmental control of *arnB* transcription in *Klebsiella* that is distinct from the pattern found in *E. coli* and *Salmonella.* Future work may uncover additional signals that also control these networks *in vivo* or in other environments.

## Materials and Methods

### Strains and Plasmids

Strains, plasmids, and primers are listed in Tables 2, 3, and 4, respectively. *E. coli* strains were derived from *E. coli* K-12 strain MG1655 (CGSC #7740), *S.* Typhimurium strains were derived from *S.* Typhimurium 14028s, and *K. pneumoniae* strains were derived from *K. pneumoniae* MGH 78578 (ATCC 700721).

**Table 2.**
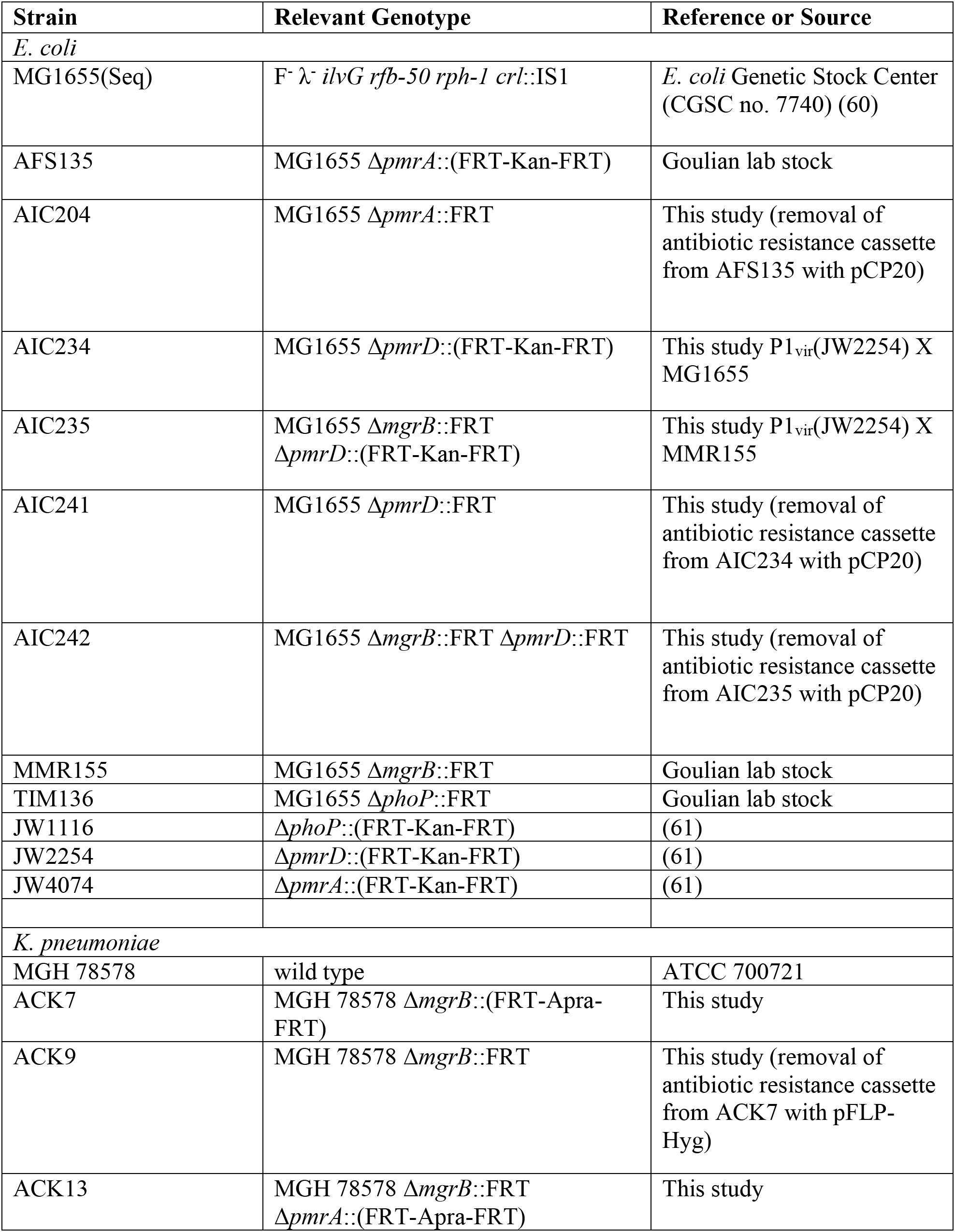
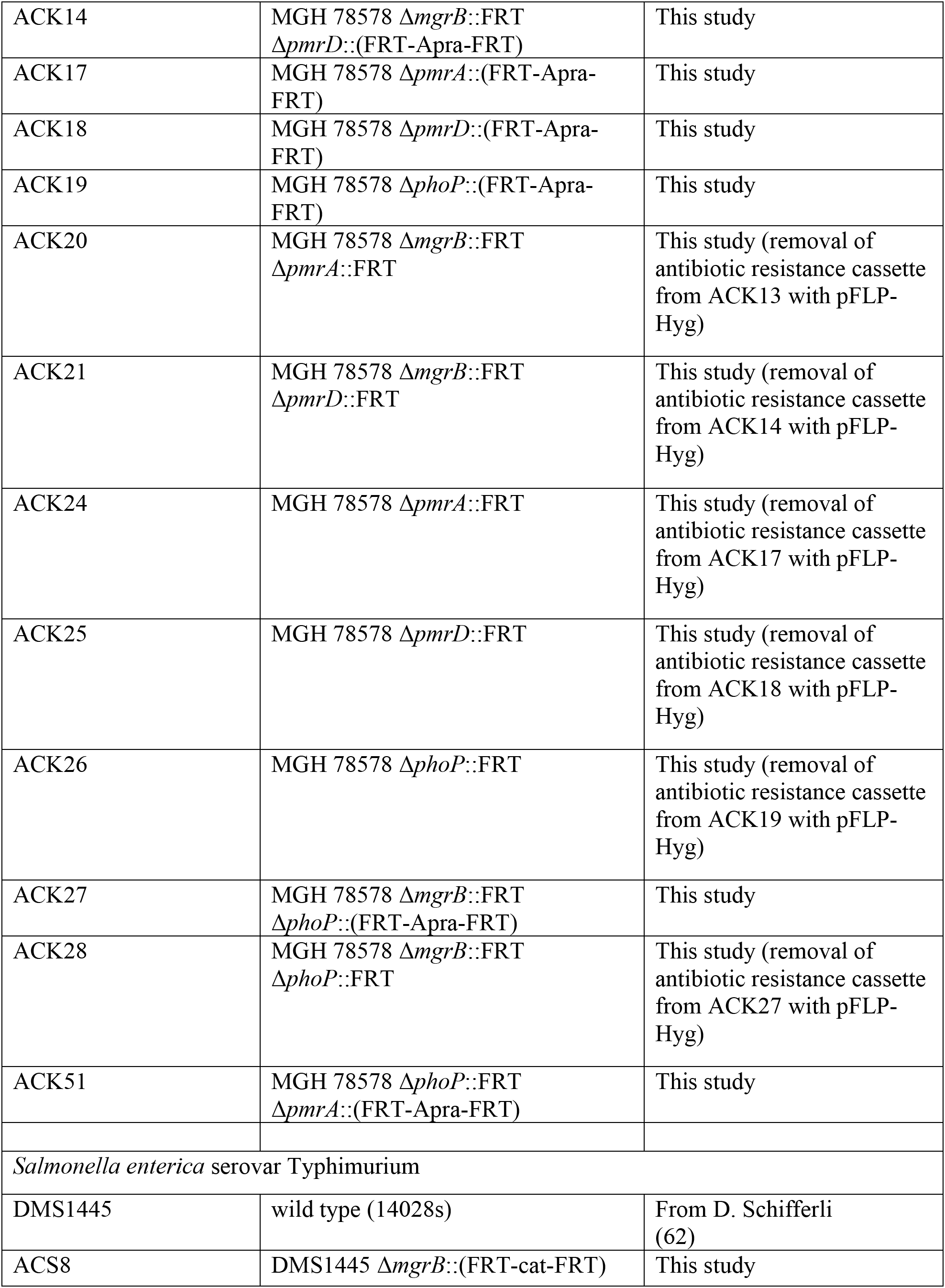
Strains used in this study

**Table 3.** Plasmids used in this study

| Plasmid | Genotype | Reference or Source |
| --- | --- | --- |
| pAC25 | Contains FRT-Apra-FRT, oriR6K | This study |
| pACBSR-Hyg | Contains arabinose-inducible lambda red recombinase genes, Hyg<sup>R</sup>, p15A origin | (57) |
| pCP20 | λcI857(ts) *repA101*(ts) *oriR101*<br>λpR-FLP *bla cat* | (54) |
| pFLP-Hyg | p15A replicon, contains FLP recombinase, Hyg<sup>R</sup> | (57) |
| pKD3 | Contains FRT-cat-FRT, oriR6K | (55) |
| pKD13 | Contains FRT-Kan-FRT, oriR6K | (55) |
| pKD46 | Contains arabinose-inducible lambda red recombinase genes, Amp<sup>R</sup>, pSC101 origin | (55) |
| pMDIAI | Contains FRT-Apra-FRT, ColE1 origin | (63) |
| pP<sub>arnB</sub>-gfp | Reporter for *E. coli* *arnB* promoter | (64) |
| pWKS130 | Control plasmid without *gfp*, Kan<sup>R</sup> | (65) |

**Table 4.**
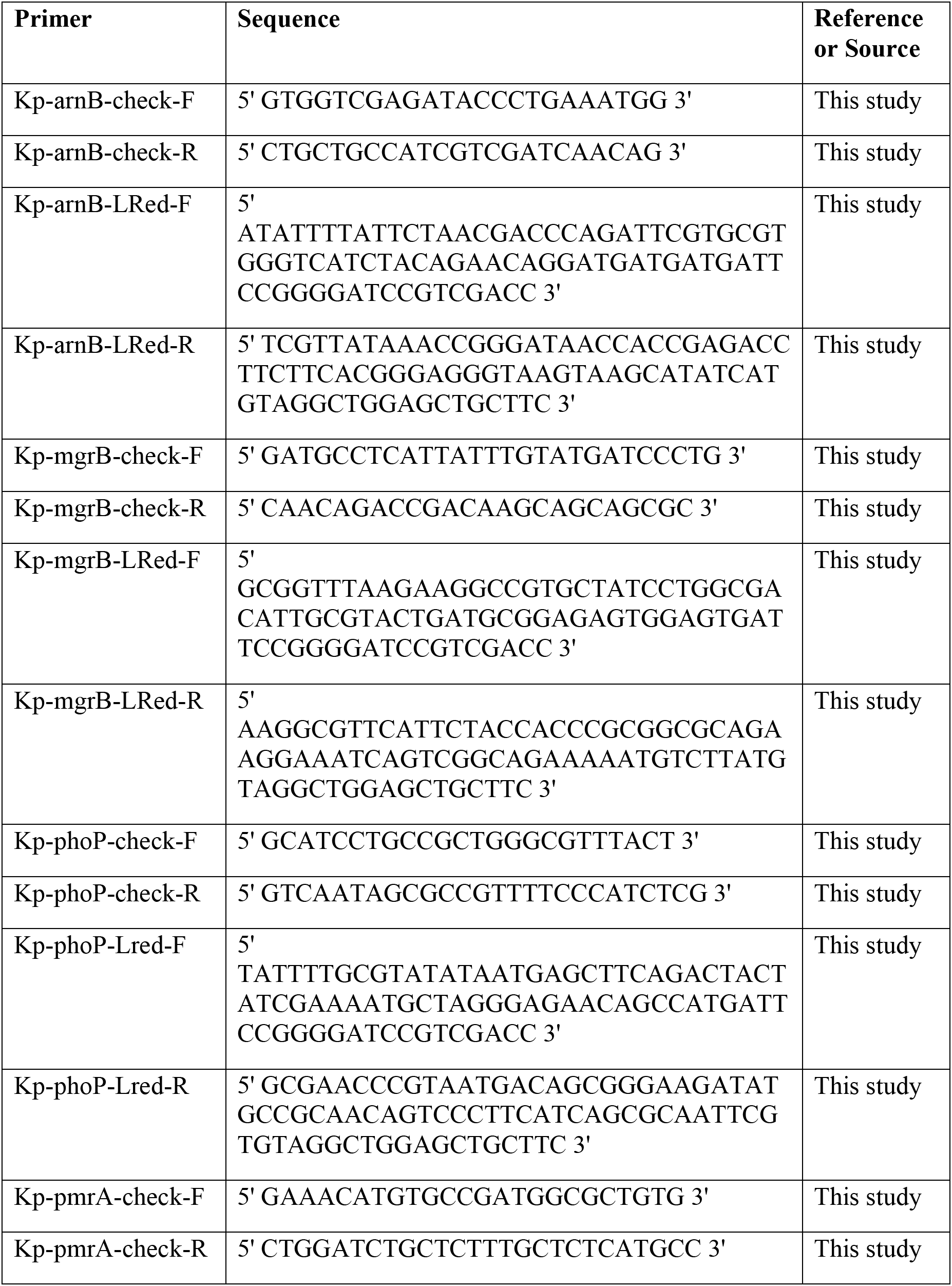
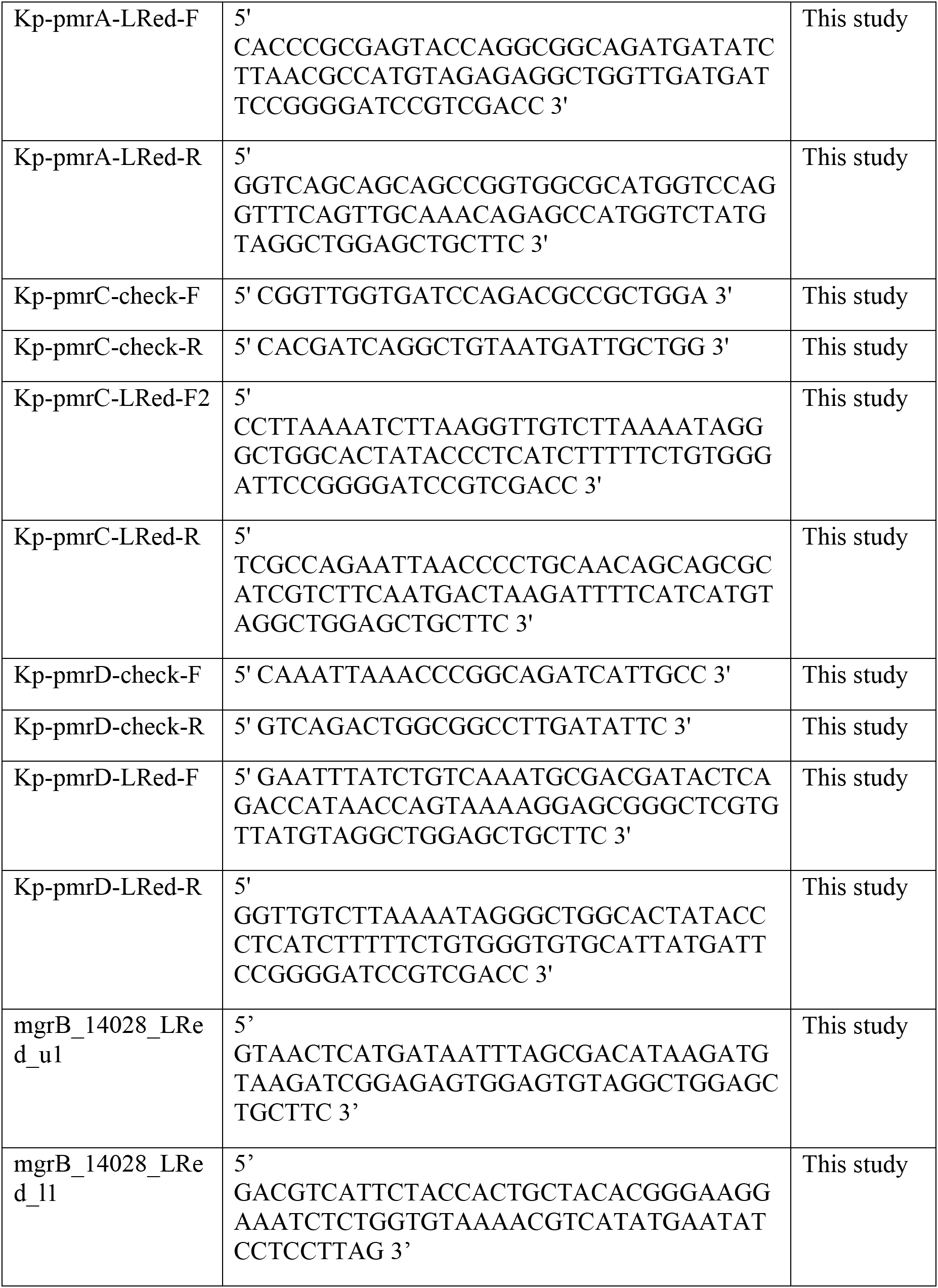
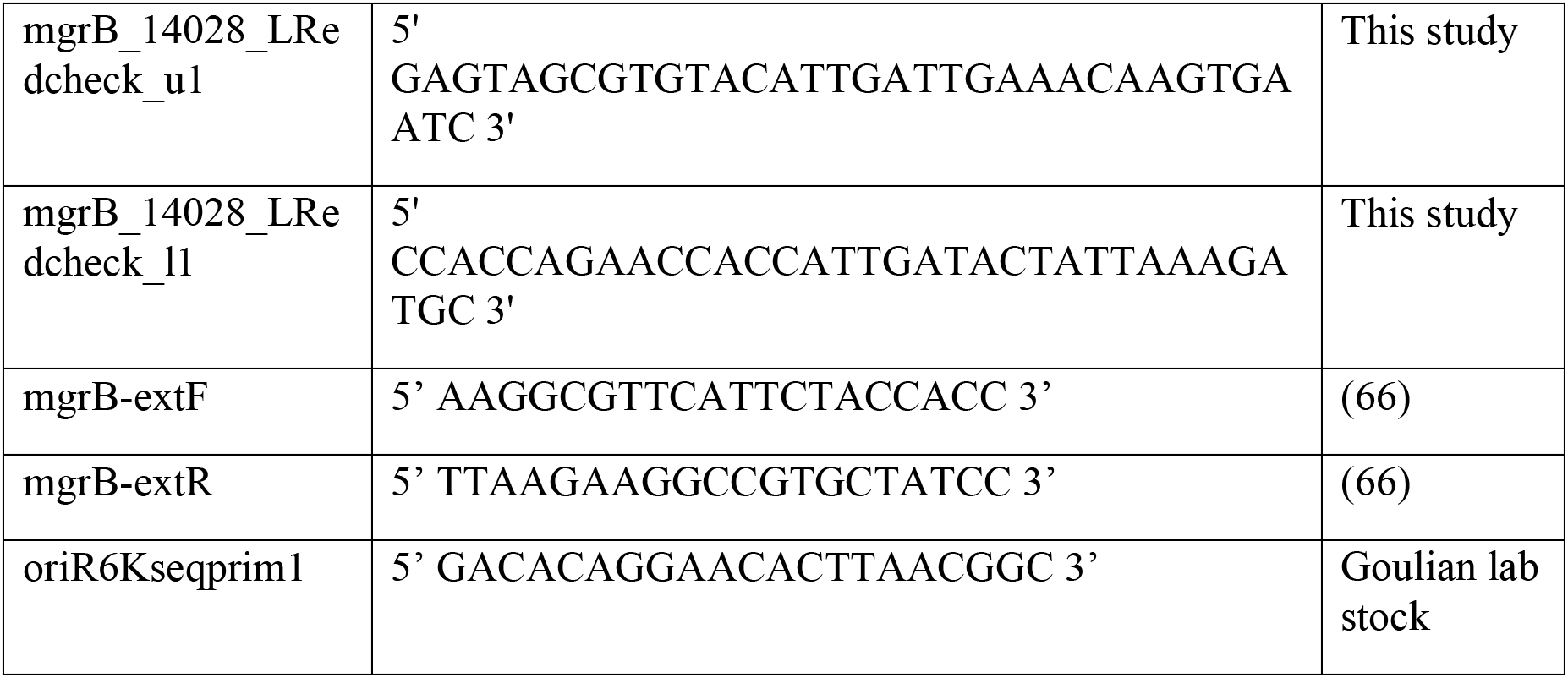
Primers used in this study

### Media and growth conditions

Cultures were grown aerobically at 37 °C in LB medium, low salt LB medium (5 g/L yeast extract, 10 g/L tryptone, and 5 g/L NaCl adjusted to pH 8.0 with NaOH), Mueller-Hinton broth (MHB), cation-adjusted MHB (MHB with 20-25 mg/L Ca^+2^ and 10-12.5 mg/L Mg^+2^), or in N-minimal medium (0.1 M Bis-Tris, pH 7.5 or 5.8, 5 mM KCl, 7.5 mM (NH_4_)_2_SO_4_, 0.5 mM K_2_SO_4_, and 1 mM KH_2_PO_4_) containing 0.2% glucose, 0.1% casamino acids, and the indicated concentrations of MgCl_2_ and FeSO_4_. Low salt LB was only used for growing cultures with hygromycin. LB (Miller) agar was used for growth on solid medium. Phosphate-buffered saline (PBS) consisted of 137 mM NaCl, 2.7 mM KCl, 10 mM Na_2_HPO_4_, and 2 mM KH_2_PO_4_. The antibiotics streptomycin, kanamycin, ampicillin, chloramphenicol, hygromycin, and apramycin were used at concentrations of 100 μg/mL, 25 μg/mL, 100 μg/mL, 25 μg/mL, 100 μg/mL, and 50 μg/mL.

### Strain Construction

#### E. coli

Gene deletions in *E. coli* were transferred to strains by transduction with P1_vir_ (53). Antibiotic resistance cassettes flanked by FLP recombinase target (FRT) sites were removed using pCP20 (54).

#### *S.* Typhimurium

The *mgrB* gene in *S.* Typhimurium was deleted and replaced with a FRT-*cat*-FRT DNA segment by lambda red recombineering using the helper plasmid pKD46 and the plasmid pKD3 as the template for FRT-*cat*-FRT, as described previously (55). The deletion was transferred to a fresh 14028s background by transduction with P22 phage. Transductants were streaked on green indicator plates (10 g/L tryptone, 5 g/L yeast extract, 5 g/L NaCl, 15 g/L agar, 0.25% glucose, 0.5% K_2_HPO_4_, 0.00125% methylene blue, and 0.0025% sodium fluorescein) to identify phage-free colonies (56).

#### K. pneumoniae

Gene deletions in *K. pneumoniae* were constructed by replacing the gene of interest with an apramycin resistance cassette FRT-*apra*-FRT by recombineering using pACBSR-Hyg as described in (57). Plasmid pMDIAI was used as the template for the FRT-*apra*-FRT segment for the Δ*mgrB* strain. For all other deletions, plasmid pAC25 was used as the template for the FRT-*apra*-FRT segment. Antibiotic resistance cassettes flanked by FLP recombinase target (FRT) sites were removed using pFLP-Hyg.

### Plasmid Construction

Plasmid pAC25 was constructed by replacing the kanamycin resistance gene in pKD13 with the apramycin resistance gene using FLP recombinase expressed from pFLP-Hyg. The plasmid was confirmed by sequencing using the primer oriR6Kseqprim1.

### Fluorescence quantification

Strains were grown overnight in N-minimal medium pH 7.5 with 1mM MgCl_2_ to saturation at 37 °C. Cultures were diluted 1:1000 into fresh medium at the indicated pH and with the indicated concentrations of MgCl_2_ and FeSO_4_ in plastic culture tubes and grown to an OD_600_ ~ 0.3-0.4. Streptomycin was added to a final concentration of 100 μg/mL, and cultures were put on ice. GFP fluorescence was measured in a fluorometer (Photon Technology International Quantamaster). Fluorescence was then divided by OD_600_ to normalize by cell density, and background fluorescence from non-fluorescent cells, measured from the strain MG1655 containing pWKS130, a control plasmid without *gfp*, was subtracted from each measurement.

### Minimum inhibitory concentration assay

Minimum inhibitory concentrations (MIC’s) of polymyxin B were determined by a broth microdilution assay as described previously (58). Briefly, cells were grown overnight in MHB to saturation at 37 °C. Cultures were diluted 1:1000 into cation-adjusted MHB and grown to OD_600_ ~ 0.3-0.4. Cells were diluted 1:150 in cation-adjusted MHB, and 50 μL was added to a 96-well microtiter plate containing two-fold dilutions of polymyxin B. The microtiter plate was incubated on a shaker at 37 °C overnight, and the MIC was determined based on turbidity.

### Polymyxin B susceptibility assay

Susceptibility assays were performed essentially as described previously (41). Strains were grown overnight in N-minimal medium pH 7.5 with 1 mM MgCl_2_ to saturation at 37 °C. Cultures were diluted 1:1000 into fresh medium at the indicated pH and concentrations of MgCl_2_ and FeSO_4_ in plastic culture tubes and grown to an OD_600_ ~ 0.3-0.4. Cultures were diluted 1:200 into Eppendorf tubes (pre-incubated at 37 °C for at least one hour) containing 1 mL PBS + 1 mM MgCl_2_ with and without 1 μg/mL polymyxin B and incubated at 37 °C for one hour. Serial dilutions in PBS were then plated in triplicate on LB + 40 mM MgCl_2_ and incubated overnight at 30 °C. Percent survival was calculated by dividing the number of colony forming units (CFUs) for samples treated with polymyxin by the corresponding CFUs of the untreated samples and multiplying by 100.

### Frequency of spontaneous polymyxin B resistance

Strains were grown in LB overnight at 37 °C. A volume of 50 μL was spread on LB containing 10 μg/mL PMB and incubated at 37 °C overnight. The initial number of CFUs plated was calculated by plating serial dilutions of the overnight culture in triplicate on LB agar. The frequency was calculated as the number of spontaneous polymyxin-resistant colonies divided by the initial number of CFUs plated. To determine whether resistant colonies of *K. pneumoniae* had mutations in *mgrB*, the *mgrB* locus was amplified using primers mgrB-extF and mgrB-extR. The DNA fragment that did not appear to have an insertion based on gel electrophoresis was sequenced using the primer mgrB-extR.

## Acknowledgements

This work was supported by NIH grants GM080279 (MG), AI120489 (JZ), and AI125814 (MG and JZ).

## References

1. Falagas ME, Kasiakou SK. 2005. Colistin: the revival of polymyxins for the management of multidrug-resistant gram-negative bacterial infections. Clin Infect Dis 40:1333–41.

2. Hancock RE. 1984. Alterations in outer membrane permeability. Annu Rev Microbiol 38:237–64.

3. Vaara M. 1992. Agents that increase the permeability of the outer membrane. Microbiol Rev 56:395–411.

4. Vaara M, Vaara T. 1983. Polycations as outer membrane-disorganizing agents. Antimicrob Agents Chemother 24:114–22.

5. Nikaido H. 2003. Molecular basis of bacterial outer membrane permeability revisited. Microbiol Mol Biol Rev 67:593–656.

6. Trimble MJ, Mlynarcik P, Kolar M, Hancock RE. 2016. Polymyxin: Alternative Mechanisms of Action and Resistance. Cold Spring Harb Perspect Med 6.

7. Olaitan AO, Morand S, Rolain JM. 2014. Mechanisms of polymyxin resistance: acquired and intrinsic resistance in bacteria. Front Microbiol 5:643.

8. Campos MA, Vargas MA, Regueiro V, Llompart CM, Alberti S, Bengoechea JA. 2004. Capsule polysaccharide mediates bacterial resistance to antimicrobial peptides. Infect Immun 72:7107–14.

9. Chen HD, Groisman EA. 2013. The biology of the PmrA/PmrB two-component system: the major regulator of lipopolysaccharide modifications. Annu Rev Microbiol 67:83–112.

10. Guo L, Lim KB, Gunn JS, Bainbridge B, Darveau RP, Hackett M, Miller SI. 1997. Regulation of lipid A modifications by Salmonella typhimurium virulence genes phoP-phoQ. Science 276:250–3.

11. Groisman EA. 2001. The pleiotropic two-component regulatory system PhoP-PhoQ. J Bacteriol 183:1835–42.

12. Wosten MM, Groisman EA. 1999. Molecular characterization of the PmrA regulon. J Biol Chem 274:27185–90.

13. Marchal K, De Keersmaecker S, Monsieurs P, van Boxel N, Lemmens K, Thijs G, Vanderleyden J, De Moor B. 2004. In silico identification and experimental validation of PmrAB targets in Salmonella typhimurium by regulatory motif detection. Genome Biol 5:R9.

14. Wosten MM, Kox LF, Chamnongpol S, Soncini FC, Groisman EA. 2000. A signal transduction system that responds to extracellular iron. Cell 103:113–25.

15. Soncini FC, Groisman EA. 1996. Two-component regulatory systems can interact to process multiple environmental signals. J Bacteriol 178:6796–801.

16. Gunn JS, Lim KB, Krueger J, Kim K, Guo L, Hackett M, Miller SI. 1998. PmrA-PmrB-regulated genes necessary for 4-aminoarabinose lipid A modification and polymyxin resistance. Mol Microbiol 27:1171–82.

17. Lee H, Hsu FF, Turk J, Groisman EA. 2004. The PmrA-regulated pmrC gene mediates phosphoethanolamine modification of lipid A and polymyxin resistance in Salmonella enterica. J Bacteriol 186:4124–33.

18. Gunn JS, Miller SI. 1996. PhoP-PhoQ activates transcription of pmrAB, encoding a two-component regulatory system involved in Salmonella typhimurium antimicrobial peptide resistance. J Bacteriol 178:6857–64.

19. Garcia Vescovi E, Soncini FC, Groisman EA. 1996. Mg2+ as an extracellular signal: environmental regulation of Salmonella virulence. Cell 84:165–74.

20. Bader MW, Navarre WW, Shiau W, Nikaido H, Frye JG, McClelland M, Fang FC, Miller SI. 2003. Regulation of Salmonella typhimurium virulence gene expression by cationic antimicrobial peptides. Mol Microbiol 50:219–30.

21. Bader MW, Sanowar S, Daley ME, Schneider AR, Cho U, Xu W, Klevit RE, Le Moual H, Miller SI. 2005. Recognition of antimicrobial peptides by a bacterial sensor kinase. Cell 122:461–72.

22. Kox LF, Wosten MM, Groisman EA. 2000. A small protein that mediates the activation of a two-component system by another two-component system. EMBO J 19:1861–72.

23. Gunn JS, Ryan SS, Van Velkinburgh JC, Ernst RK, Miller SI. 2000. Genetic and functional analysis of a PmrA-PmrB-regulated locus necessary for lipopolysaccharide modification, antimicrobial peptide resistance, and oral virulence of Salmonella enterica serovar typhimurium. Infect Immun 68:6139–46.

24. Kato A, Groisman EA. 2004. Connecting two-component regulatory systems by a protein that protects a response regulator from dephosphorylation by its cognate sensor. Genes Dev 18:2302–13.

25. Cheng HY, Chen YF, Peng HL. 2010. Molecular characterization of the PhoPQ-PmrD-PmrAB mediated pathway regulating polymyxin B resistance in Klebsiella pneumoniae CG43. J Biomed Sci 17:60.

26. Mitrophanov AY, Groisman EA. 2008. Signal integration in bacterial two-component regulatory systems. Genes Dev 22:2601–11.

27. Mitrophanov AY, Jewett MW, Hadley TJ, Groisman EA. 2008. Evolution and dynamics of regulatory architectures controlling polymyxin B resistance in enteric bacteria. PLoS Genet 4:e1000233.

28. Guckes KR, Breland EJ, Zhang EW, Hanks SC, Gill NK, Algood HM, Schmitz JE, Stratton CW, Hadjifrangiskou M. 2017. Signaling by two-component system noncognate partners promotes intrinsic tolerance to polymyxin B in uropathogenic Escherichia coli. Sci Signal 10.

29. Wright MS, Suzuki Y, Jones MB, Marshall SH, Rudin SD, van Duin D, Kaye K, Jacobs MR, Bonomo RA, Adams MD. 2015. Genomic and transcriptomic analyses of colistin-resistant clinical isolates of Klebsiella pneumoniae reveal multiple pathways of resistance. Antimicrob Agents Chemother 59:536–43.

30. Cheng YH, Lin TL, Lin YT, Wang JT. 2016. Amino Acid Substitutions of CrrB Responsible for Resistance to Colistin through CrrC in Klebsiella pneumoniae. Antimicrob Agents Chemother 60:3709–16.

31. Poirel L, Jayol A, Nordmann P. 2017. Polymyxins: Antibacterial Activity, Susceptibility Testing, and Resistance Mechanisms Encoded by Plasmids or Chromosomes. Clin Microbiol Rev 30:557–596.

32. Olaitan AO, Diene SM, Kempf M, Berrazeg M, Bakour S, Gupta SK, Thongmalayvong B, Akkhavong K, Somphavong S, Paboriboune P, Chaisiri K, Komalamisra C, Adelowo OO, Fagade OE, Banjo OA, Oke AJ, Adler A, Assous MV, Morand S, Raoult D, Rolain JM. 2014. Worldwide emergence of colistin resistance in Klebsiella pneumoniae from healthy humans and patients in Lao PDR, Thailand, Israel, Nigeria and France owing to inactivation of the PhoP/PhoQ regulator mgrB: an epidemiological and molecular study. Int J Antimicrob Agents 44:500–7.

33. Cannatelli A, Giani T, D’Andrea MM, Di Pilato V, Arena F, Conte V, Tryfinopoulou K, Vatopoulos A, Rossolini GM, Group CS. 2014. MgrB inactivation is a common mechanism of colistin resistance in KPC-producing Klebsiella pneumoniae of clinical origin. Antimicrob Agents Chemother 58:5696–703.

34. Lippa AM, Goulian M. 2009. Feedback inhibition in the PhoQ/PhoP signaling system by a membrane peptide. PLoS Genet 5:e1000788.

35. Roland KL, Martin LE, Esther CR, Spitznagel JK. 1993. Spontaneous pmrA mutants of Salmonella typhimurium LT2 define a new two-component regulatory system with a possible role in virulence. J Bacteriol 175:4154–64.

36. Sun S, Negrea A, Rhen M, Andersson DI. 2009. Genetic analysis of colistin resistance in Salmonella enterica serovar Typhimurium. Antimicrob Agents Chemother 53:2298–305.

37. Zhou Z, Ribeiro AA, Lin S, Cotter RJ, Miller SI, Raetz CR. 2001. Lipid A modifications in polymyxin-resistant Salmonella typhimurium: PMRA-dependent 4-amino-4-deoxy-L-arabinose, and phosphoethanolamine incorporation. J Biol Chem 276:43111–21.

38. Phan MD, Nhu NTK, Achard MES, Forde BM, Hong KW, Chong TM, Yin WF, Chan KG, West NP, Walker MJ, Paterson DL, Beatson SA, Schembri MA. 2017. Modifications in the pmrB gene are the primary mechanism for the development of chromosomally encoded resistance to polymyxins in uropathogenic Escherichia coli. J Antimicrob Chemother 72:2729–2736.

39. Quesada A, Porrero MC, Tellez S, Palomo G, Garcia M, Dominguez L. 2015. Polymorphism of genes encoding PmrAB in colistin-resistant strains of Escherichia coli and Salmonella enterica isolated from poultry and swine. J Antimicrob Chemother 70:71–4.

40. Cannatelli A, Giani T, Aiezza N, Di Pilato V, Principe L, Luzzaro F, Galeotti CL, Rossolini GM. 2017. An allelic variant of the PmrB sensor kinase responsible for colistin resistance in an Escherichia coli strain of clinical origin. Sci Rep 7:5071.

41. Winfield MD, Groisman EA. 2004. Phenotypic differences between Salmonella and Escherichia coli resulting from the disparate regulation of homologous genes. Proc Natl Acad Sci U S A 101:17162–7.

42. Chen HD, Jewett MW, Groisman EA. 2011. Ancestral genes can control the ability of horizontally acquired loci to confer new traits. PLoS Genet 7:e1002184.

43. Rubin EJ, Herrera CM, Crofts AA, Trent MS. 2015. PmrD is required for modifications to escherichia coli endotoxin that promote antimicrobial resistance. Antimicrob Agents Chemother 59:2051–61.

44. Groisman EA, Kayser J, Soncini FC. 1997. Regulation of polymyxin resistance and adaptation to low-Mg2+ environments. J Bacteriol 179:7040–5.

45. Kidd TJ, Mills G, Sa-Pessoa J, Dumigan A, Frank CG, Insua JL, Ingram R, Hobley L, Bengoechea JA. 2017. A Klebsiella pneumoniae antibiotic resistance mechanism that subdues host defences and promotes virulence. EMBO Mol Med 9:430–447.

46. Prost LR, Daley ME, Le Sage V, Bader MW, Le Moual H, Klevit RE, Miller SI. 2007. Activation of the bacterial sensor kinase PhoQ by acidic pH. Mol Cell 26:165–74.

47. Perez JC, Groisman EA. 2007. Acid pH activation of the PmrA/PmrB two-component regulatory system of Salmonella enterica. Mol Microbiol 63:283–93.

48. Alpuche Aranda CM, Swanson JA, Loomis WP, Miller SI. 1992. Salmonella typhimurium activates virulence gene transcription within acidified macrophage phagosomes. Proc Natl Acad Sci U S A 89:10079–83.

49. Miyashiro T, Goulian M. 2007. Stimulus-dependent differential regulation in the Escherichia coli PhoQ PhoP system. Proc Natl Acad Sci U S A 104:16305–10.

50. Yadavalli SS, Carey JN, Leibman RS, Chen AI, Stern AM, Roggiani M, Lippa AM, Goulian M. 2016. Antimicrobial peptides trigger a division block in Escherichia coli through stimulation of a signalling system. Nat Commun 7:12340.

51. Mangan S, Alon U. 2003. Structure and function of the feed-forward loop network motif. Proc Natl Acad Sci U S A 100:11980–5.

52. Mangan S, Zaslaver A, Alon U. 2003. The coherent feedforward loop serves as a sign-sensitive delay element in transcription networks. J Mol Biol 334:197–204.

53. Miller JH. 1992. A Short Course in Bacterial Genetics. Cold Spring Harbor Laboratory.

54. Cherepanov PP, Wackernagel W. 1995. Gene disruption in Escherichia coli: TcR and KmR cassettes with the option of Flp-catalyzed excision of the antibiotic-resistance determinant. Gene 158:9–14.

55. Datsenko KA, Wanner BL. 2000. One-step inactivation of chromosomal genes in Escherichia coli K-12 using PCR products. Proc Natl Acad Sci U S A 97:6640–5.

56. Lopez CA, Winter SE, Rivera-Chavez F, Xavier MN, Poon V, Nuccio SP, Tsolis RM, Baumler AJ. 2012. Phage-mediated acquisition of a type III secreted effector protein boosts growth of salmonella by nitrate respiration. MBio 3.

57. Huang TW, Lam I, Chang HY, Tsai SF, Palsson BO, Charusanti P. 2014. Capsule deletion via a lambda-Red knockout system perturbs biofilm formation and fimbriae expression in Klebsiella pneumoniae MGH 78578. BMC Res Notes 7:13.

58. CLSI. 2012. Methods for Dilution Antimicrobial Susceptibility Tests for Bacteria that Grow Aerobically; Approved Standard-Ninth Edition. Institute CaLS, Wayne, PA.

59. Kato A, Latifi T, Groisman EA. 2003. Closing the loop: the PmrA/PmrB two-component system negatively controls expression of its posttranscriptional activator PmrD. Proc Natl Acad Sci U S A 100:4706–11.

60. Freddolino PL, Amini S, Tavazoie S. 2012. Newly identified genetic variations in common Escherichia coli MG1655 stock cultures. J Bacteriol 194:303–6.

61. Baba T, Ara T, Hasegawa M, Takai Y, Okumura Y, Baba M, Datsenko KA, Tomita M, Wanner BL, Mori H. 2006. Construction of Escherichia coli K-12 in-frame, single-gene knockout mutants: the Keio collection. Mol Syst Biol 2:2006 0008.

62. Miller SI, Mekalanos JJ. 1990. Constitutive expression of the phoP regulon attenuates Salmonella virulence and survival within macrophages. J Bacteriol 172:2485–90.

63. Yang J, Sun B, Huang H, Jiang Y, Diao L, Chen B, Xu C, Wang X, Liu J, Jiang W, Yang S. 2014. High-efficiency scarless genetic modification in Escherichia coli by using lambda red recombination and I-SceI cleavage. Appl Environ Microbiol 80:3826–34.

64. Zaslaver A, Bren A, Ronen M, Itzkovitz S, Kikoin I, Shavit S, Liebermeister W, Surette MG, Alon U. 2006. A comprehensive library of fluorescent transcriptional reporters for Escherichia coli. Nat Methods 3:623–8.

65. Wang RF, Kushner SR. 1991. Construction of versatile low-copy-number vectors for cloning, sequencing and gene expression in Escherichia coli. Gene 100:195–9.

66. Cannatelli A, D’Andrea MM, Giani T, Di Pilato V, Arena F, Ambretti S, Gaibani P, Rossolini GM. 2013. In vivo emergence of colistin resistance in Klebsiella pneumoniae producing KPC-type carbapenemases mediated by insertional inactivation of the PhoQ/PhoP mgrB regulator. Antimicrob Agents Chemother 57:5521–6.

